# Structural changes in secondary, but not primary, sensory cortex in individuals with congenital olfactory sensory loss

**DOI:** 10.1101/760793

**Authors:** Moa G. Peter, Gustav Mårtensson, Elbrich M. Postma, Love Engström Nordin, Eric Westman, Sanne Boesveldt, Johan N. Lundström

## Abstract

Individuals with congenital sensory loss usually demonstrate altered brain morphology in areas associated with early processing of the lost sense. Here, we aimed to establish whether this also applies to individuals born without a sense of smell (congenital anosmia) by comparing cortical morphology between 33 individuals with isolated congenital anosmia and matched controls. We detected no structural alterations in the primary olfactory (piriform) cortex. However, individuals with anosmia demonstrated gray matter volume atrophy in bilateral olfactory sulci, explained by decreased cortical area, curvature, and sulcus depth. They further demonstrated increased gray matter volume and cortical thickness in the medial orbital gyri; regions closely associated with olfactory processing, sensory integration, and value-coding. Our results suggest that a lifelong absence of sensory input does not necessarily lead to morphological alterations in primary sensory cortex and extend previous findings with divergent morphological alterations in bilateral orbitofrontal cortex, indicating influences of different plastic processes.

## INTRODUCTION

The notion that the human brain is plastic and undergoes morphological as well as functional alterations in response to changes in experienced demands is widely accepted (Buonomano and Merzenich 1998; Lindenberger et al. 2017). One of the more drastic changes in demands on the brain is undoubtedly the loss of a sensory modality, comprising a complete lack of input from the lost sense combined with altered demands on the remaining senses. Indeed, visual sensory loss has repeatedly been linked to both structural and functional cerebral reorganizations with often profound changes in regions normally focused on the processing of the lost sense, often in primary sensory cortex (Frasnelli et al. 2011; Merabet and Pascual-Leone 2010; for reviews see Bavelier and Neville 2002). In contrast, little is known about the potential plastic effects of complete olfactory sensory loss (anosmia) and especially whether individuals with lifelong (congenital) anosmia display cerebral reorganization.

Although the brain exhibits plasticity throughout life, it is strongest early in life when even brief periods of sensory deprivation can make it difficult, if not impossible, to gain normal abilities even if the sensory loss is reversed and normal sensory input established (Collignon et al. 2015; Hyvärinen et al. 1981; Guerreiro et al. 2016; Wiesel and Hubel 1965). Thus, in comparison to individuals who have gone through normal sensory development, individuals with a congenital or very early acquired complete sensory loss would be expected to demonstrate pronounced patterns of cerebral reorganization. In addition, studying individuals with an isolated congenital sensory loss (i.e., a congenital sensory loss not related to additional symptoms) has the advantage of isolating plastic effects of the sensory deprivation, whereas individuals with an acquired sensory loss constitute a much more heterogeneous group where variability in the age at which the sense was lost, the duration of the sensory loss, the direct cause of sensory loss (as in the case of traumatic brain injury, which might in itself cause reorganization of the brain), and perceptual abilities before sensory loss likely affect sensory loss-related cerebral reorganization (Noppeney et al. 2005; Voss and Zatorre 2012; Jiang et al. 2015). Individuals with isolated congenital sensory loss hence constitute a good model for increasing our understanding of the adaptiveness the human brain possesses.

Both congenital and acquired visual sensory loss have repeatedly been linked to atrophy in form of decreased gray matter volume in areas related to visual processing, specifically within primary visual cortex (Ptito et al. 2008; Bridge et al. 2009; Jiang et al. 2015; Noppeney et al. 2005; Pan et al. 2007). The gray matter volume decreases are, however, associated with divergent underlying morphology: the atrophy in congenital blindness is accompanied by a thickening of the visual cortex (Park et al. 2009; Hasson et al. 2016; Bridge et al. 2009; Jiang et al. 2009), whereas the atrophy in individuals with acquired sensory loss has been linked to either a cortical thinning or a lack of cortical thickness alterations (Park et al. 2009; Jiang et al. 2009; Voss and Zatorre 2012). The fact that divergent underlying morphology may cause similar volumetric results emphasize the importance of not solely relying on one measure of morphological changes.

Despite the established link between olfactory ability and the morphology of olfactory cortical structures such as the olfactory bulb, piriform (commonly referred to as primary olfactory) cortex, and orbitofrontal cortex (OFC; commonly referred to as secondary olfactory cortex), which generally indicates a positive correlation between volume and ability (Seubert, Freiherr, Frasnelli, et al. 2013; Hummel et al. 2015; Frasnelli et al. 2010), the only consistent finding in the study of cerebral morphological reorganization in individuals with isolated congenital anosmia (ICA) is the absence, or hypoplasia, of the olfactory bulbs and olfactory tracts, accompanied by a significant decrease in olfactory sulcus depth (Abolmaali et al. 2002; Huart et al. 2011; Yousem et al. 1996). There are, to the best of our knowledge, only two studies that have investigated whether individuals with ICA display cortical alterations beyond the olfactory bulb and sulcus in a statistical manner: one by Frasnelli and colleagues (2013) and one by Karstensen and colleagues (2018). Both studies indicate that individuals with ICA have increased gray matter volume in primary olfactory cortex (piriform cortex), albeit in opposite hemispheres. Morphological alterations within primary sensory cortex are in agreement with the visual loss literature, but in contrast to the atrophy in congenitally blind, ICA seems to be linked to a gray matter volume increase. In addition to primary olfactory cortex, alterations within secondary olfactory (orbitofrontal) cortex are reported by both Karstensen and Frasnelli: the former finds a volume decrease around the left posterior olfactory sulcus, and although the latter finds no evidence in support of volumetric alterations in OFC, an increase in cortical thickness in bilateral anteromedial OFC is reported. Albeit the overlap between the two studies is small, the failure of exact replication does not suggest inaccurate results, but indicates that the sample sizes are likely not large enough to gain sufficient statistical power for stable results. This is not a unique problem. Sample sizes in studies of congenital sensory loss, independent of sensory modality explored, are typically small (undoubtedly a consequence of the scarcity of these conditions) and it is uncommon to include more than 20 patients.

In the present study, we aimed to determine whether cortical morphological alterations beyond the olfactory bulb and olfactory sulcus depth are present in individuals with congenital anosmia. Structural magnetic resonance imaging data was collected for 34 individuals with ICA (analyzed for 33, see methods) and 34 normosmic controls, matched in terms of age, sex, and education. First, we determined potential group differences in whole-brain gray matter volume using voxel-based morphometry (VBM) for voxel-wise comparisons. Thereafter, to assess possible underlying mechanisms of the VBM results, we determined potential differences in cortical thickness, surface area, and curvature between individuals with ICA and normosmic controls. Based on past findings indicating that both congenital blindness and ICA leads to alterations in cortical thickness and/or gray matter volume within early processing areas of the lost sense, we hypothesized that individuals with ICA would demonstrate an increase in gray matter volume within piriform cortex and OFC.

## MATERIALS AND METHODS

### Participants

A total of 68 participants were enrolled in the study: 34 individuals with isolated congenital anosmia (ICA, congenital anosmia unrelated to specific genetic disorders, such as Kallmann syndrome) and 34 controls, matched in terms of sex, age, and educational level (Table 1). Inclusion criteria for the ICA group was a lifelong lack of olfactory perception with no known underlying condition causing the anosmia and for the control group a self-proclaimed functional sense of smell (subsequently tested). In addition, 24 out of the 34 individuals with ICA had received a diagnosis from a physician at a previous occasion. Participants were recruited and tested at two different sites: 46 participants (23 ICA) in Stockholm, Sweden, and 22 (11 ICA) in Wageningen, the Netherlands; the matched control was always tested at the same site as the individual with ICA. One individual from the ICA group was removed from analysis after visual inspection of the images due to abnormal anatomy, leaving a final sample of 33 individuals with ICA and 34 controls. All participants provided written informed consent and the study was approved by the ethical review boards in both Sweden and in the Netherlands.

**Table 1.** Descriptive statistics per experimental group.

|  | ICA (n = 33) | Control (n = 34) |
| --- | --- | --- |
| Age (years) | 34.2 (12.9) | 34 (12.1) |
| Female | 21 | 22 |
| Education (years) | 14.1 (2.6) | 14.1 (1.7) |
| TDI | 10.9 (2.3) | 35.4 (3.8) |
| *Threshold* | 1.2 (0.5) | 8.7 (3.2) |
| *Discrimination* | 4.9 (1.6) | 13.5 (1.6) |
| *Identification* | 4.8 (1.5) | 13.4 (1.3) |
| Left olfactory sulcus depth (mm) | 4.9 (3.3) | 8.6 (2.3) |
| Right olfactory sulcus depth (mm) | 5.8 (3.6) | 8.8 (2.3) |

Values presented as mean (standard deviation). ICA = Isolated congenital anosmia, TDI = combined score from the Sniffin' Sticks olfactory sub-tests.

### Procedure

#### Olfactory screening

Olfactory function was assessed with the full Sniffin’ Sticks olfactory test (Burghart, Wedel, Germany), a standardized test consisting of three subtests with individual scores: odor detection threshold (T), odor quality discrimination (D), and 4-alternative cued odor quality identification (I), together yielding the combined TDI-score. Mean TDI scores on the Sniffin’ Stick olfactory performance test were 10.9 (SD = 2.3, range: 7-15) and 35.4 (SD = 3.8, range: 28.5-42.5) for ICA and controls, respectively (Table 1), with all individuals in the ICA group demonstrating olfactory scores below the limit for functional anosmia (TDI cut-off at 16.0; Oleszkiewicz et al. 2019).

#### Image acquisition

Imaging data was acquired on two 3T Siemens Magnetom MR scanners (Siemens Healthcare, Erlangen, Germany). At the Swedish site, data was acquired on a Prisma scanner using a 20-channel head coil and in the Netherlands, a Verio scanner with a 32-channel head coil. The scanning sequence protocols were identical at both sites.

T1-weighted images with whole-brain coverage were acquired using an MP-RAGE sequence (TR = 1900 ms, TI = 900 ms, TE = 2.52 ms, flip angle = 9°, voxel size = 1 mm3, 176 slices, FoV = 256 mm) and T2-weighted images with whole-brain coverage were acquired using a SPACE sequence (TR = 3200 ms, TE = 405 ms, voxel size = 1 mm3, 224 slices, FoV = 256 mm). Additional functional imaging data, not reported here, were collected after the anatomical images in the same testing session.

### Image processing

All images were visually inspected to ensure high image quality with absence of unwanted artifacts before inclusion. Three types of measures were extracted. Cortical volumetric measures to assess potential effects of ICA on grey matter volume; surface based measures to assess potential mechanisms of volumetric results; olfactory sulcus depth to support clinical diagnose of ICA and to replicate past findings of differences in olfactory sulcus depth between ICA and control individuals (Yousem et al. 1996; Abolmaali et al. 2002; Huart et al. 2011).

#### Volumetric measures

Voxel-based morphometry (VBM, Ashburner and Friston 2000) analysis was done using SPM12 (Wellcome Trust Centre for Neuroimaging, UCL; http://www.fil.ion.ucl.ac.uk/spm) in MATLAB 2016a (The MathWorks, Inc., Natick, Massachusetts, USA). T1-weighted images were segmented into gray matter, white matter, and cerebrospinal fluid in native space using unified segmentation (Ashburner and Friston 2005). The three tissues were used to compute total intracranial volume for each participant. The gray and white matter were further used as input to a diffeomorphic image registration algorithm to improve inter-subject alignment (DARTEL, Ashburner 2007). DARTEL implements an iterative process in which gray and white matter from all subjects are aligned, creating an increasingly accurate average template for inter-subject alignment. The template and individual flow fields from DARTEL were then used to spatially normalize gray matter images to MNI space with 12 parameter affine transformations. The normalized gray matter images were modulated with the Jacobian determinant of the deformation fields and smoothed with a 6 mm full-width at half maximum isotropic Gaussian kernel. The relatively small kernel was chosen to optimize the discovery of group differences in piriform cortex, a small area at the junction between frontal and temporal cortex.

#### Surface based measures

Surface based measures were used to calculate cortical thickness, surface area, and underlying white matter curvature. The T1-weighted images were processed using FreeSurfer ver. 6.0 (Dale et al. 1999; http://surfer.nmr.mgh.harvard.edu/). The processing pipeline is described in detail elsewhere (Fischl and Dale 2000), but, in short, the image processing included removal of non-brain tissue, Talairach transformation, segmentation of subcortical white matter and gray matter, intensity normalization, tessellation of the boundary between gray and white matter, automated topology correction, and surface deformation to find the gray/white matter boundary and the gray matter/cerebrospinal fluid boundary. Cortical thickness, area, and mean curvature of the gray/white matter boundary was calculated at each vertex point on the tessellated surface in native space. Thereafter, individual surfaces were aligned to an average template and smoothed (5 mm full-width at half maximum surface based Gaussian kernel chosen to optimize analysis in piriform cortex) to enable statistical comparisons between the subject groups. FreeSurfer data was pre-processed through the HiveDB database system (Muehlboeck et al. 2014).

#### Olfactory sulcus depth

Two independent raters, blind to participant group, measured the depth of the olfactory sulci in the plane of the posterior tangent through the eyeballs, according to the method proposed by Huart and colleagues (2011). In short, on the first slice towards the posterior where the eyeballs were no longer seen on the T2-weighted MR images, a line measuring the depth of the olfactory sulcus was drawn, using the Multi-Image Analysis GUI (Mango version 4.0.1; Research Imaging Institute, UTHSCSA http://ric.uthscsa.edu/mango). There was a high agreement between the two raters’ measurements (intraclass correlation coefficient of .88). Sulci defined as outliers (sulci for which the two initial raters’ measurements differed more than 2 standard deviations from the mean difference; in total one sulcus in three individuals and both sulci in three individuals) were assessed by an additional observer. Olfactory sulcus depth was defined as the obtained mean measure, independent of the number of raters.

### Statistical analysis

#### Volumetric measures

To compensate for individual differences in intracranial volume when comparing gray matter volume between groups, the pre-processed, normalized gray matter images were proportionally scaled with total intracranial volume. Furthermore, the images were masked with a threshold of 0.15 to avoid inclusion of non-gray matter voxels. Voxel-wise differences in gray matter volume between groups were estimated with independent sample t-tests with age, sex, and scanning site as nuisance covariates. The tests were corrected for multiple comparisons in a whole-brain analysis with a family-wise error (FWE) corrected significance level of *p* < .05.

#### Surface based measures

To investigate whether potential volumetric differences can be explained by differences in cortical thickness, surface area, or curvature, vertex-wise values were compared between groups using a general linear model with age, sex, and scanning site as nuisance covariates. Correction for multiple comparisons was done based on a false discovery rate (FDR) of < .05 and group differences surviving this threshold were defined as significant. If no vertex survived this threshold, a more liberal threshold of *p* < .001, uncorrected, was used to explore the full extent of potential differences; highlighted in text when used.

#### Olfactory sulcus depth

To assess differences between the two groups in olfactory sulcus depth, a repeated measures analysis of variance (rmANOVA; within subject factor Hemisphere, between subject factor Group) was used; effect size estimates are reported as partial eta-squared (*η*_*p*_^*2*^). Potential group differences in olfactory sulcus depth were thereafter investigated for the right and left olfactory sulcus separately using Welch’s t-test. Effects size estimates are given by Cohen’s d.

## RESULTS

### Altered morphology within the orbitofrontal cortex in congenital anosmia

To explore volumetric alterations, voxel-wise tests of group differences in gray matter volume were performed in a VBM analysis. Five clusters passed the FWE-corrected significance level of *p* < .05. All of the significant clusters were located in the OFC: two demonstrating gray matter atrophy in the ICA group around the bilateral olfactory sulci, two clusters demonstrating a gray matter volume increase in the ICA group centered around the bilateral medial orbital gyri (the latter located anterolateral to the aforementioned two clusters of gray matter atrophy), and an additional small cluster of increased volume in the right posterior orbital gyrus (Figure 1, Table 2).

**Table 2.** Results from volumetric measures.

| Region | Hemisphere | x<sup>a</sup> | y<sup>a</sup> | z<sup>a</sup> | Cluster size<sup>b</sup> | t<sup>c</sup> | p<sup>d</sup> |
| --- | --- | --- | --- | --- | --- | --- | --- |
| *ICA < Control* |  |  |  |  |  |  |  |
| Olfactory sulcus<sup>1</sup> | L | -12 | 21 | -16 | 2185 | 10.5 | < .001 |
| Olfactory sulcus<sup>2</sup> | R | 13 | 19 | -13 | 2135 | 9.7 | < .001 |
| *ICA > Control* |  |  |  |  |  |  |  |
| Medial orbital gyrus<sup>3</sup> | L | -21 | 39 | -21 | 532 | 6.7 | .001 |
| Medial orbital gyrus<sup>4</sup> | R | 17 | 39 | -14 | 244 | 6.0 | .006 |
| Posterior orbital sulcus,<br>middle branch | R | 22 | 24 | -11 | 8 | 5.5 | .04 |

**Figure 1.**
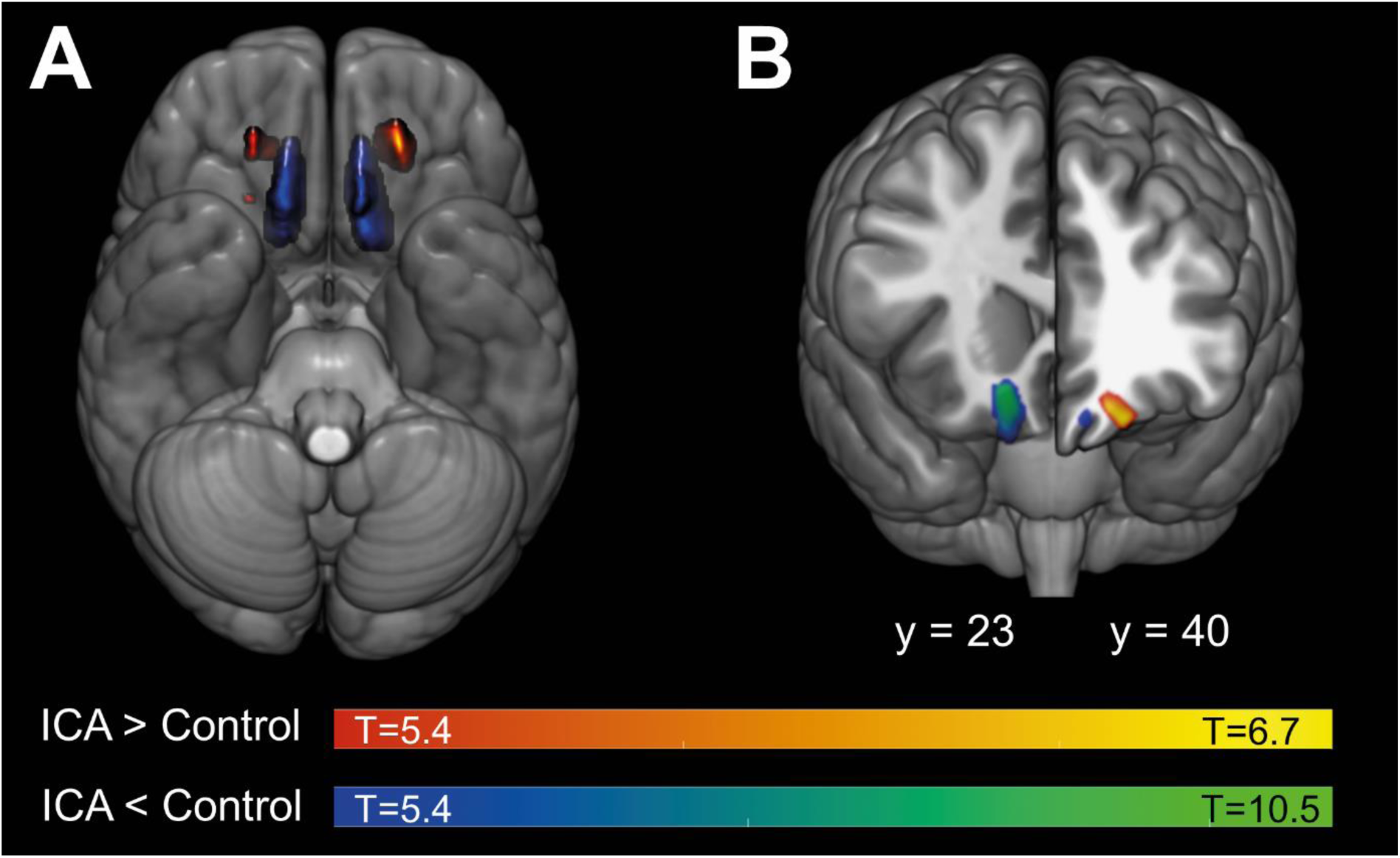
Groups differences in gray matter volume. Gray matter volume differences displayed on MNI152-template. Individuals with isolated congenital anosmia (ICA) demonstrate gray matter volume atrophy in the bilateral olfactory sulci as well as gray matter volume increases in bilateral medial orbital gyri. **A** Results displayed on an inferior view of a transparent brain to reveal full extent of clusters. **B** Results displayed on an anterior view with coronal section at y=23 (right hemisphere) and y=40 (left hemisphere). All results are displayed at the statistical level of p < .05, FWE-corrected.

In opposition to our initial hypothesis, no significant group differences in gray matter volume within piriform (primary olfactory) cortex were demonstrated at an FWE-corrected significance level of *p* < .05. Given our specific hypothesis, two additional steps were taken. First, to confirm that the lack of group differences were not simply an effect of an excessively conservative statistical threshold, the significance level was altered to *p* < .01, uncorrected. Even at this liberal statistical threshold, no significant group differences in either left or right piriform cortex could be detected (Supplementary Figure S1). Second, to investigate whether there is statistical evidence in support of the lack of group differences, mean gray matter volume within the piriform cortex was extracted based on the region of interest from Zhou and colleagues (2019) using MarsBaR toolbox (http://marsbar.sourceforge.net/). Equivalence testing between groups was performed using the two one-sided tests procedure for equivalence testing using the TOSTER R package (Lakens 2017). The gray matter volume within both right and left piriform cortex were statistically equivalent between groups within the equivalence bounds we have 80% power to detect (corresponding to an effect size of *d* ≥ .6; the test yielding the highest *p*-value in the right and left hemisphere, respectively: *t*(62.7) = 2.16, *p* = .017; *t*(61.8) = 2.33, *p* = .012), indicating that it is unlikely that there are real statistical differences between groups.

To investigate whether the gray matter volume alterations demonstrated within the ICA group could be further explained by alterations in cortical thickness, surface area, or curvature, these measures were compared vertex-wise between groups. The cortical thickness analysis demonstrated a thickening of the left medial orbital gyrus and anterior olfactory sulcus in the ICA group (Figure 2A, Table 3); however, no group differences in cortical thickness were found in the right hemisphere when using the designated statistical cut-off (FDR < .05). When applying a more liberal threshold (*p* < .001, uncorrected), clusters similar to the ones in the left hemisphere were revealed in the right hemisphere, i.e., a cortical thickening around the olfactory sulcus in the ICA group (Supplementary Figure S2). In addition to altered gray matter volume and cortical thickness in the OFC, the ICA group further demonstrated decreased surface area along the bilateral olfactory sulci (Figure 2B, Table 3) accompanied by decreased curvature around the same orbitofrontal areas: the bilateral olfactory sulci and medial orbital gyri (Figure 2C, Table 3), all at an FDR threshold of < .05. Furthermore, a small cluster of increased curvature was demonstrated in the left superior temporal sulcus (Table 3, Supplementary Figure S3).

**Table 3.** Results from surface based measures.

| Region | Hemisphere | x<sup>a</sup> | y<sup>a</sup> | z<sup>a</sup> | Cluster size<sup>b</sup> | p<sup>c</sup> |
| --- | --- | --- | --- | --- | --- | --- |
| **Cortical Thickness** |  |  |  |  |  |  |
| *ICA > Control* |  |  |  |  |  |  |
| Medial orbital gyrus<sup>5</sup> | L | -15 | 35 | -22 | 171 | 10<sup>-10</sup> |
| Posteromedial orbital lobule | L | -16 | 8 | -16 | 12 | 10<sup>-4.4</sup> |
| **Area** |  |  |  |  |  |  |
| *ICA < Control* |  |  |  |  |  |  |
| Olfactory sulcus<sup>6</sup> | R | 14 | 16 | -13 | 343 | 10<sup>-8.4</sup> |
| Olfactory sulcus<sup>7</sup> | L | -14 | 9 | -16 | 251 | 10<sup>-9</sup> |
| **Curvature** |  |  |  |  |  |  |
| *ICA < Control* |  |  |  |  |  |  |
| Olfactory sulcus<sup>8</sup> | L | -12 | 37 | -20 | 77 | 10<sup>-12</sup> |
| Olfactory sulcus<sup>9</sup> | R | 16 | 10 | -15 | 56 | 10<sup>-6.6</sup> |
| Medial orbital gyrus<sup>10</sup> | L | -14 | 42 | -22 | 35 | 10<sup>-5.9</sup> |
| Olfactory sulcus<sup>11</sup> | R | 15 | 33 | -19 | 34 | 10<sup>-6.3</sup> |
| Superior temporal sulcus | L | -55 | -43 | 7 | 22 | 10<sup>-4.7</sup> |
| Olfactory sulcus<sup>12</sup> | L | -16 | 10 | -16 | 16 | 10<sup>-5.8</sup> |
| Medial orbital gyrus<sup>13</sup> | R | 14 | 25 | -25 | 15 | 10<sup>-5.6</sup> |
| Medial/posterior orbital gyrus | L | -17 | 15 | -22 | 9 | 10<sup>-5.2</sup> |
| Medial/ posterior orbital gyrus | R | 16 | 17 | -23 | 2 | 10<sup>-4.2</sup> |

**Figure 2.**
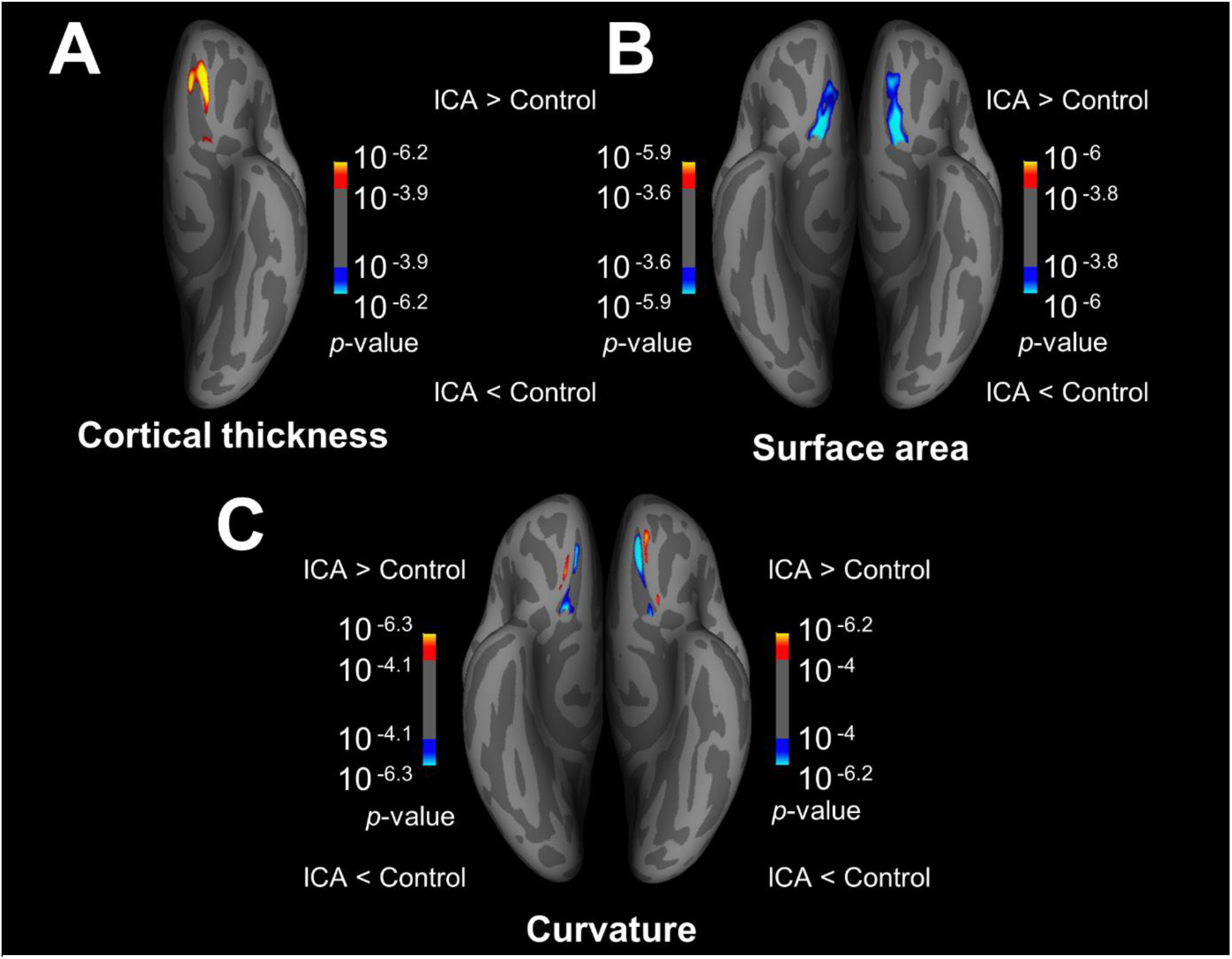
Group differences in surface based measures. Results are displayed on inflated brain, inferior view. Displayed results are corrected for multiple comparisons based on a false discovery rate (FDR) < .05. **A** Differences in cortical thickness. Individuals with ICA demonstrate a significant increase in cortical thickness in the left medial orbital gyrus/olfactory sulcus. No significant group differences were demonstrated in the right hemisphere at FDR < .05, for visual comparison of the right hemisphere results thresholded at p < .001 uncorrected, see supplementary Figure S2. **B** Group differences in surface area demonstrate a significantly decreased area for individuals with ICA along the bilateral olfactory sulci. **C** Group differences in curvature. Significant differences demonstrated in bilateral olfactory sulcus and bilateral medial orbital gyrus; all indicating a lower absolute value of curvature for individuals with ICA, i.e., larger radii of curves (FreeSurfer’s convention of curvature is negative for gyri and positive for sulci, warm colors indicate ICA > Control).

### Decreased olfactory sulcus depth in individuals with congenital anosmia

Finally, we assessed potential differences between individuals with ICA and normosmic controls in olfactory sulcus depth were investigated with a rmANOVA. Individuals with ICA demonstrated significantly more shallow olfactory sulci as compared to controls, main effect of Group: F(1,65) = 23.7, *p* < .001, *η*_*p*_^*2*^ = .267; ICA mean = 5.3 mm, SD = 3.4 mm; Control mean = 8.7 mm, SD = 2.3 mm. Furthermore, a difference of depth in the right, as compared to left, olfactory sulcus was demonstrated, main effect of Hemisphere: *F*(1,65) = 9.0, *p* = .004, *η*_*p*_^*2*^ = .120; however, no significant interaction between Group and Side was present, *F*(1,65) = 2.8, *p* = .098, *η*_*p*_^*2*^ = .042. The decrease in olfactory sulcus depth for the ICA group was demonstrated for the left olfactory sulcus, *t*(57.6) = 5.28, *p* < .0001, *d* = 1.3, as well as for the right olfactory sulcus, *t*(54.3) = 4.11, *p* < .001, *d* = 1.01; Figure 3 and Table 1.

**Figure 3.**
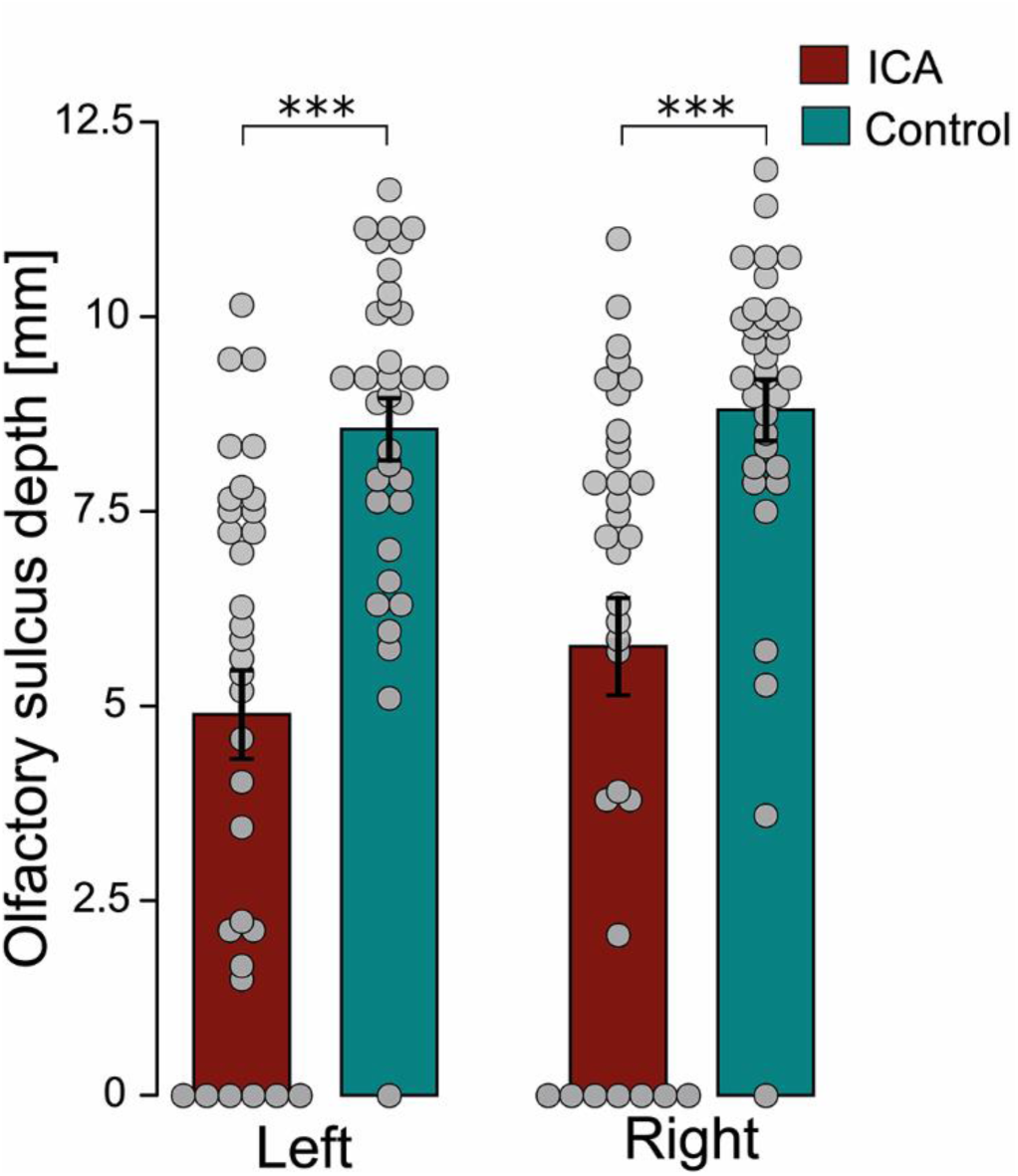
Group differences in olfactory sulcus depth. Mean olfactory sulcus depth (error bars indicate standard error) overlaid with individual values and separated by hemisphere and group. Individuals with Isolated Congenital Anosmia (ICA) demonstrate a significant decrease in bilateral olfactory sulci compared to matched controls. *** indicating p < .001.

### Relationships between olfactory sulcus depth and morphology results

Both volumetric and surface based analysis revealed structural group differences in bilateral medial OFC, mainly in, or in close proximity to, the olfactory sulci. Based on these results, we assessed whether the volumetric and surface based group differences were linked to the group difference in olfactory sulcus depth by means of Pearson’s correlation coefficient analyses. For each individual, men gray matter volume was computed for each of the four large significant clustersSimilarly, mean cortical thickness was computed for the large significant cluster in the left hemisphere, mean surface area for the two significant clusters, and mean curvature was computed in the three largest significant cluster in left OFC and the three largest significant clusters in right OFC (all clusters used for correlation analysis marked in Table 2 and Table 3). Pearson’s correlation coefficients were computed between each of these structural measures and the olfactory sulcus depth in the corresponding hemisphere, with Bonferroni correction applied for number of tests of each type of structural measure (4 for volume, 2 for area, 6 for curvature). To avoid that the large demonstrated differences between groups would influence the correlations, the Pearson’s correlation coefficients were computed within each group separately.

Both the ICA and control group demonstrated significant negative correlation between olfactory sulcus depth and cortical thickness, as well as between olfactory sulcus depth and curvature, in the left medial orbital gyrus (Table 4). Furthermore, the curvature in the neighboring cluster in the left anterior olfactory sulcus correlated with olfactory sulcus depth in the control (but not ICA) group, whereas corresponding cluster in the right hemisphere (anterior olfactory sulcus) correlated with olfactory sulcus depth in the ICA group. Significant correlations between sulcus depth and surface area in the olfactory sulci were demonstrated, with exception for the left sulcus in the ICA group. Additionally, the gray matter volume cluster along the right olfactory sulcus correlated with the olfactory sulcus depth in the ICA, but not control, group. No other correlations between gray matter volume and olfactory sulcus depth were significant (Table 4).

**Table 4.** Correlations between listed clusters and olfactory sulcus depth in corresponding hemisphere.

| Region | Hemisphere | ICA |  | Control |  |
| --- | --- | --- | --- | --- | --- |
|  |  | *r* | *p* | *r* | *p* |
| VBM |  |  |  |  |  |
| Olfactory sulcus<sup>1</sup> | L | .28 | .109 | .31 | .072 |
| Olfactory sulcus<sup>2</sup> | R | .52 | .002 ** | .25 | .16 |
| Medial orbital gyrus<sup>3</sup> | L | .12 | .491 | -.24 | .166 |
| Medial orbital gyrus<sup>4</sup> | R | .06 | .726 | -.31 | .066 |
| Cortical thickness |  |  |  |  |  |
| Medial orbital gyrus<sup>5</sup> | L | -.51 | .003 ** | -.59 | 2.7·10<sup>-4</sup> *** |
| Area |  |  |  |  |  |
| Olfactory sulcus<sup>6</sup> | R | .43 | .013 * | .39 | .023 * |
| Olfactory sulcus<sup>7</sup> | L | .28 | .111 | .42 | .013 * |
| Curvature |  |  |  |  |  |
| Olfactory sulcus<sup>8</sup> | L | .24 | .172 | .50 | .003 * |
| Olfactory sulcus<sup>9</sup> | R | .36 | .040 | .00 | .981 |
| Medial orbital gyrus<sup>10</sup> | L | -.5 | .003 * | -.63 | 6.3·10<sup>-5</sup> *** |
| Olfactory sulcus<sup>11</sup> | R | .65 | 3.9·10<sup>-5</sup> *** | .12 | .485 |
| Olfactory sulcus<sup>12</sup> | L | .3 | .087 | .24 | .168 |
| Medial orbital gyrus<sup>13</sup> | R | -.32 | .072 | -.06 | .745 |

## DISCUSSION

We can here demonstrate that individuals with isolated congenital anosmia (ICA) have a significantly altered structure in bilateral medial orbitofrontal cortex (OFC), a multimodal region often referred to as secondary olfactory cortex, but in contrast to our hypothesis, show no evidence of alterations within primary olfactory cortex (piriform cortex). Specifically, individuals with ICA demonstrated structural alterations, compared to normosmic individuals, in and around the olfactory sulci, encompassing gray matter volume atrophy stretching along the olfactory sulci towards the medial orbital gyri, likely linked to the decreased sulcus depth, surface area, and curvature of the olfactory sulci. Moreover, gray matter volume and cortical thickness increases were demonstrated in the medial orbital gyri of individuals with ICA. These divergent results in neighboring cortical areas indicate that the cerebral plasticity related to a life-long olfactory sensory deprivation cannot be explained by a simple morphological mechanism applied to all olfactory-related cerebral areas, but rather, that the plasticity is area- or network-dependent.

The cortical structural reorganization demonstrated by individuals with ICA were restricted to secondary olfactory (medial orbitofrontal) cortex, without any indications of structural alterations in the primary olfactory (piriform) cortex. This lack of reorganization in primary sensory cortex is in disagreement with the literature on congenital visual sensory loss (Park et al. 2009; Jiang et al. 2009; Hasson et al. 2016; Bridge et al. 2009), as well as previous studies on individuals with congenital anosmia (Frasnelli et al. 2013; Karstensen et al. 2018). Although the two previous studies investigating cortical morphology in individuals with congenital anosmia indicate increased cortical thickness and increased gray matter volume within the piriform cortex, their results are unilateral and in opposite hemispheres. Albeit our results are contradictory to both the aforementioned results and our own hypothesis, the lack of morphological group differences in the piriform cortex reported here are arguably reliable as they are based on considerably larger samples than the two previous studies. In addition, neither volumetric nor the individual surface based measures detected alterations in piriform cortex; measures that did, however, indicate clear morphological reorganization in medial orbitofrontal areas. It could be argued that ICA does lead to structural alterations in piriform cortex but that the effects are modest, thus limiting the possibility of discovery. Speaking against this argument is both the lack of observable differences also when applying a liberal statistical threshold uncorrected for multiple comparisons, and the fact that equivalence tests indicated that the gray matter volume within the piriform cortices were equal between groups, within the bounds detectable with the (in this context) large sample size. Nonetheless, the piriform cortex is a small structure with complex folding and to firmly establish that a lifelong lack of olfactory perception does not alter its structure, studies with higher spatial resolution should be conducted to get a more precise estimate of the piriform morphology in this subject group. Moreover, it should be noted that a lack of morphological changes in primary olfactory cortex does not necessarily indicate that no functional reorganization due to absent olfactory sensory input has occurred.

Even though piriform cortex is commonly referred to as primary olfactory cortex, either by itself or along with the other recipients of olfactory bulb projections, it can be argued that the piriform cortex should rather be labeled as an odor association cortex due to its involvement in more complex processing, beyond what is commonly associated with primary sensory areas (Gottfried 2010). Among others, anterior piriform cortex has been associated with both attention to, and working memory for, odors (Zelano et al. 2005; Zelano et al. 2009). The fact that piriform cortex performs functions more similar to association cortex in other sensory systems could potentially explain discrepancies in results obtained in congenitally blind and individuals with ICA when it comes to morphological alterations in primary sensory cortex. The lack of input to the piriform cortex from the olfactory bulb might in ICA individuals be compensated for by input from downstream regions, such as the amygdala, OFC, and insula. However, what role the piriform cortex have in individuals with ICA needs to be established before mechanistic explanations to the lack of morphological alterations within the piriform cortex can be elucidated. Nonetheless, although debating what does constitute primary olfactory cortex is beyond the scope of this article, we can conclude that no evidence of ICA-dependent morphological alterations in any of the cerebral areas (beyond the olfactory bulb) commonly mentioned as candidates were demonstrated, even at a very liberal statistical threshold within a comparably large sample.

Our results demonstrate that the OFC of individuals with ICA has a distinctly different structure than that of normosmic individuals. Although it is evident that this cortical reorganization is strongly related to the congenital sensory loss, the results are in their nature insufficient to conclude how these anomalies emerged. One hypothesis is that individuals with ICA are born with an abnormality in the cortical folding of the medial OFC, manifested as a cortical flattening around the olfactory sulci and medial orbital gyri, linked to the absence, or hypoplasia, of the olfactory bulbs and tracts. The olfactory bulbs and tracts reside in the olfactory sulci and the projection of the olfactory tracts is suggested to play an important role in the formation and deepening of the olfactory sulci during early development (Turetsky et al. 2009; Abolmaali et al. 2002; Huart et al. 2011). A congenital abnormal cortical folding could explain the decreased olfactory sulcus depth, decreased area, and decreased curvature in olfactory sulci and medial orbital gyri, as well as the gray matter volume atrophy in the olfactory sulci, as a natural consequence of the decreased surface area. However, a decrease in cortical surface in the pericalcarine sulci (visual cortex) in congenitally blind individuals has been described by Park and colleagues (Park et al. 2009), suggesting that the decreased olfactory sulcus depth in individuals with ICA might not be exclusively caused by a congenital abnormal anatomy linked to small or absent olfactory bulbs and tracts, but, similar to the blind, be a consequence of atypical development caused by a lack of sensory input. In addition, although the theory of congenital abnormal cortical folding in individuals with ICA provides a straightforward explanation of the gray matter volume atrophy and corresponding surface based measures, it cannot fully explain the gray matter volume increases in the medial orbital gyri. By taking the proposed function of these cortical areas into account, other possible explanations of the cortical reorganization emerge.

The medial OFC is alongside the piriform cortex considered vital for olfactory processing (Lundström et al. 2011; Seubert, Freiherr, Djordjevic, et al. 2013; Gottfried 2006) and the cortical regions in which the individuals with ICA demonstrated cortical thickness and gray matter volume increases show a substantial overlap with areas repeatedly implicated in olfactory processing (Seubert, Freiherr, Djordjevic, et al. 2013; for reviews see Gottfried and Zald 2005). Considering these orbitofrontal areas as predominantly olfactory, the cortical increases displayed by individuals with ICA are presumably caused by the fact that these areas have never received olfactory input. In contrast, individuals with acquired olfactory sensory loss tend to demonstrate cortical volume decreases in the OFC (Yao et al. 2017; Bitter et al. 2010). Although these morphological consequences of olfactory sensory loss might seem contradictory, similar results have repeatedly been demonstrated in blind individuals, with a thickening of visual cortex of congenitally blind and a thinning of those with acquired blindness (Park et al. 2009; Voss and Zatorre 2012). In addition, the demonstrated cortical volume and thickness increases reported here are an extension of Frasnelli and colleagues’ (2013) report of increased cortical thickness in the OFC of individuals with ICA. Cortical increases as a result of a lack of sensory processing might seem counter intuitive and the mechanisms behind these morphological alterations are not yet fully understood. One of the main theories proposed to explain the cortical increases in congenitally blind individuals (Park et al. 2009; Jiang et al. 2009), as well as in individuals with ICA (Frasnelli et al. 2013), is that the complete lack of sensory input to a cortical area normally devoted to the processing of the lost sensory modality alters normal postnatal development. Specifically, the absent input results in a lack of the typical sensory input-driven synaptic pruning of redundant connections. The exact mechanisms of these results need to be addressed using animal models.

Although the OFC undoubtedly is important for olfactory processing (Li et al. 2010), it is a much more multimodal area than the visual areas showing cortical thickness increases in congenitally blind. Generally, the key functions of the OFC is often put in the context of value encoding, including, but by no means limited to the olfactory domain (Gottfried and Zelano 2011; Kringelbach and Rolls 2004; van den Bosch et al. 2014), with direct stimulation of human OFC eliciting not only olfactory but also somatosensory, gustatory, and emotional experiences (Fox et al. 2018). Specifically, the OFC is relevant for the processing and integration of information from different sensory modalities, receiving olfactory, gustatory, somatosensory, visceral, visual, and auditory input (Kringelbach and Rolls 2004; Rolls 2005; Ongür and Price 2000), and the medial OFC has been suggested to be of high importance for food consumption (Rolls 2005; Price 2008). If we consider the medial OFC as a multimodal rather than predominantly olfactory processing area, it can be argued that increases in gray matter volume and cortical thickness are not, as previously argued, caused by a lack of synaptic pruning due to absent olfactory input but, to the contrary, related to increased (compensatory) processing of the non-olfactory functions of the OFC. This argument is in line with data demonstrating a positive relation between cortical thickness in visual cortex and performance on auditory tasks in blind individuals (Voss and Zatorre 2012). To find support for either theory, the specific functions of the orbitofrontal areas demonstrating structural reorganization in individuals with ICA need to be further investigated, preferably using functional neuroimaging methods, to delineate whether the structural reorganization in individuals with ICA is accompanied by altered function and connectivity.

In addition to the clear reorganization in the orbitofrontal cortex, confirmed with both volumetric and surface based methods, individuals with ICA demonstrated a small but statistically significant cluster of altered curvature in the left superior temporal sulcus. The superior temporal sulcus is, along with the intraparietal sulcus and the prefrontal cortex, one of the classical cortical multisensory integration areas (Ghazanfar and Schroeder 2006; Stein and Stanford 2008; Regenbogen et al. 2017). Although the group difference in the superior temporal sulcus is restricted to the curvature measure, it is noteworthy that a recent study demonstrated that individuals with ICA showed enhanced audio-visual integration performance; an effect that was interpreted as compensatory abilities due to the sensory loss (Peter et al. 2019). Whether the deviating anatomy in the superior temporal sulcus demonstrated here is in fact related to the compensatory processing of multisensory stimuli in individuals with ICA previously reported should be investigated using functional neuroimaging techniques.

In line with previous studies, we could further demonstrate that individuals with ICA have a significantly decreased olfactory sulcus depth, as compared to matched normosmic individuals (Huart et al. 2011; Abolmaali et al. 2002; Yousem et al. 1996; Levy et al. 2013; Karstensen et al. 2018). The shallow olfactory sulcus depth in individuals with ICA is thought to be related to the complete lack of, or hypoplasia of, olfactory bulbs, often accompanied by small or absent olfactory tracts (Yousem et al. 1996; Abolmaali et al. 2002); structures located in the olfactory sulci. The olfactory sulcus depth has been suggested to be an indicator of congenital anosmia (Abolmaali et al. 2002; Huart et al. 2011), an idea supported by the fact that a difference in olfactory bulb size, but not in olfactory sulcus depth, is found between individuals with acquired anosmia and controls (Rombaux et al. 2010). It is, however, important to note that even though there are significant group differences between individuals with ICA and normosmic individuals, the use of olfactory sulcus depth of a certain value as a tool to diagnose olfactory dysfunction does not allow for an unambiguous diagnose, as there is overlap between groups in sulcus depth.

The restricted spatial distribution of the cortical alterations in individuals with ICA, combined with the established decrease in olfactory sulcus depth, begs the question whether there is a direct relationship between the olfactory sulcus depth and surrounding cortical morphology. Although the correlation analyses of olfactory sulcus depth and surface area indicate that a more shallow olfactory sulcus is associated with a decreased surface area in said sulcus, and strong support for a relation between the sulcus depth and both cortical thickness and curvature in the left medial orbital gyrus was demonstrated, the other correlation measures yielded mainly non-significant results. This ambiguity could partly be related to the fact that the sulcus depth is measured at one specific location, and the underlying morphology is likely more strongly related to areas in close proximity to that location. Nonetheless, the fact that significant correlations could be demonstrated between olfactory sulcus depth and all types of cortical measures could be interpreted as support for the hypothesis of abnormal cortical folding. However, this is speculative and needs to be validated by evidence from developmental studies.

In conclusion, the current study demonstrates substantial cortical reorganization beyond the olfactory bulb and olfactory sulcus depth in individuals with ICA. Strikingly, no structural reorganization within primary olfactory (piriform) cortex is demonstrated. The lack of structural alterations in primary sensory areas indicate that the cortical reorganization mechanisms at play when a sensory system is absent is sensory modality dependent. All structural alterations, except for a small region in the superior temporal sulcus, are restricted to the secondary olfactory (orbitofrontal) cortex, and indicate areas of atrophy as well as increases in individuals born without the olfactory sense. These alterations could potentially be due to a combination of congenital abnormal morphology and a sensory loss-dependent plastic reorganization occurring during development. These clear findings of morphological changes motivate future studies of their functional and behavioral relevance.

## ACKNOWLEDGEMENTS

This work was supported by the Knut and Alice Wallenberg Foundation (KAW 2018.0152 to J.N.L.) and the Swedish Research Council (2014-1346 to J.N.L.).

