## Supplementary Figure S1, S2, S3 for "Structural changes in secondary, but not primary, sensory cortex in individuals with congenital olfactory sensory loss"

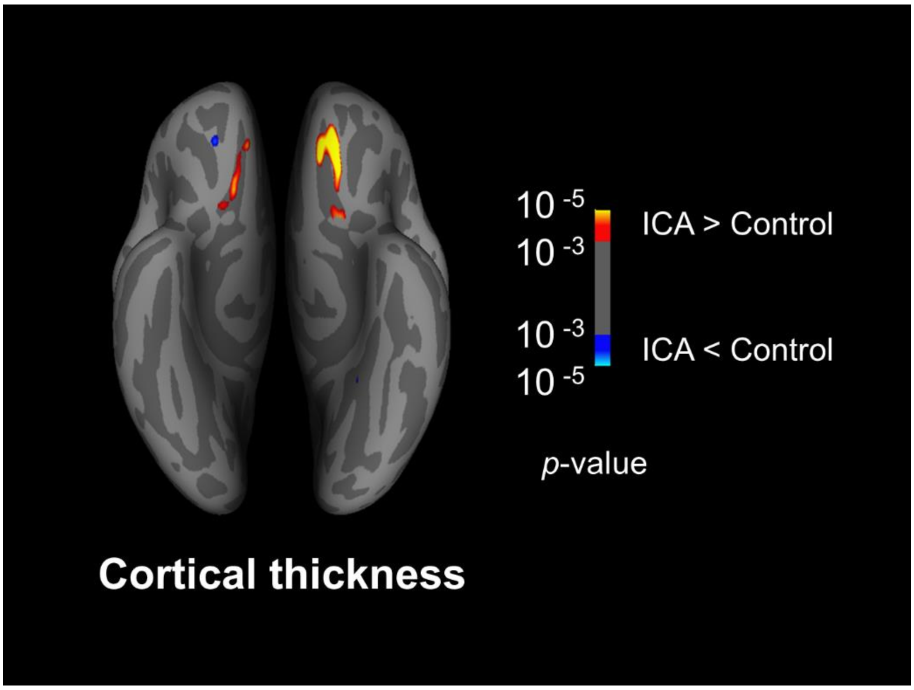


***Figure S1*** *Group differences in cortical thickness. Results are displayed on an inflated brain, inferior view, p < .001. When using a more liberal statistical threshold than the initial FDR-corrected, individuals with ICA demonstrate a similar pattern of cortical thickening in the right hemisphere as in the left.*

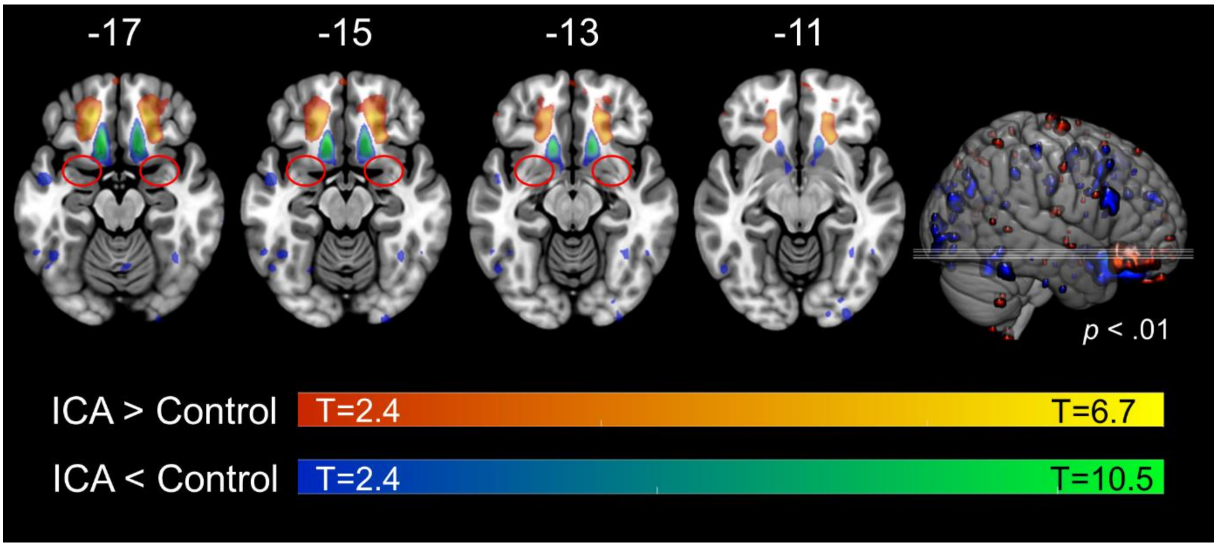


***Figure S2 – Group differences in gray matter volume.*** *No group differences in gray matter volume in piriform cortex at the liberal statistical threshold of p < .01 (uncorrected). Numbers in white above axial slices indicate z-coordinate in MNI-space and the red circles indicate the position of piriform cortex. Slice positions displayed on brain to the right.*

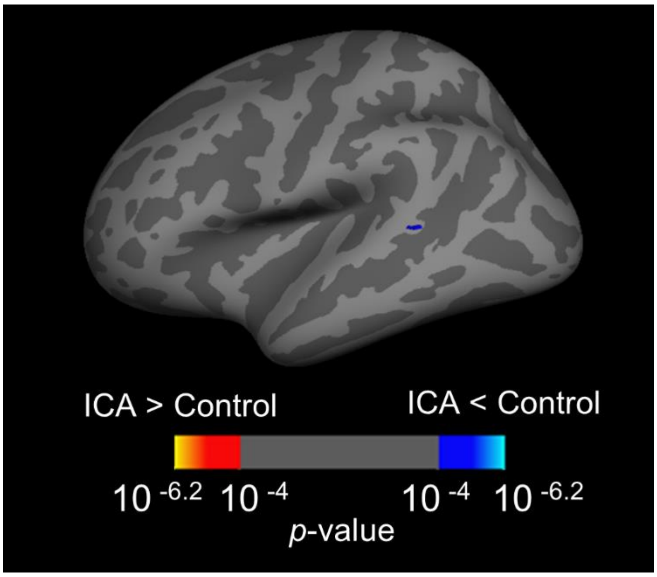


***Figure S3 - Group differences in curvature.*** *Results are displayed on inflated brain, lateral view of the left hemisphere. Individuals with isolated congenital anosmia display a cluster of decreased curvature in the superior temporal sulcus. Results thresholded at FDR < .05.*
